# Evolutionary dynamics of Gag-restricted prion-like domains in LTR retroelements

**DOI:** 10.64898/2026.11.24.769039

**Authors:** Cameron DeBellefeuille, Tomasz Zajkowski

## Abstract

**Background:** Prions are proteins that can self-template, giving rise to non-Mendelian inheritance. They are functionally diverse and have been found in all domains of life and even beyond cellular life within viruses, a distribution that suggests an ancient evolutionary origin. Testing that origin has been difficult, as prions are identified by composition, such as an enrichment of Q/N residues, and are carried by rapidly changing disordered domains that are difficult to study phylogenetically. Long terminal repeat (LTR) retroelements, mobile genetic elements that replicate through an RNA intermediate and include both retrotransposons and retroviruses, are well suited to investigate prion evolution. Prion-like domains (PrLDs) occur in their Gag proteins, while their conserved Pol proteins allow phylogenetic reconstruction independent of the domains themselves.

**Results:** Here, we screened 2,636 proteins from mobile genetic elements in the GyDB and identified 26 PrLDs. All 26 occurred in Gag, the only one of 74 protein categories to carry them, and this association remained significant when proteins were matched for intrinsic disorder, indicating that PrLDs are not a by-product of the high disorder of Gag. Gag PrLDs were found mainly in the C-terminal half of the protein. Their composition matched that of validated yeast and archaeal prion domains in glutamine and asparagine enrichment and charge depletion, with elevated aromatic content in common with the archaeal domains and an unusually high proline content. Across a phylogeny of 258 elements built from Pol, PrLD presence was clustered rather than randomly distributed, the root was reconstructed as PrLD-positive, and losses outpaced gains by roughly eighteenfold.

**Conclusions:** Our findings establish PrLDs as a general feature of Gag across LTR retroelements, reveal a proline-rich composition that sets them apart from characterised prion domains, and trace their presence to the ancestor of these elements. Their retention across three superfamilies despite frequent loss points to maintenance rather than repeated acquisition. By mapping a compositionally defined domain onto a phylogeny built from conserved proteins, this work allows the gain and loss of prion-like sequence to be reconstructed and provides 26 candidates for experimental testing.

## Background

Prions were first described as a novel class of infectious agent, distinct from any known bacterial, viral or eukaryotic pathogen [1], and were for decades regarded simply as misfolded proteins that cause disease [2]. The best-known example is bovine spongiform encephalopathy, which left 177 people and over four million cattle dead [3] and was eventually traced to a misfolded form of an endogenous protein, designated the Prion Protein (PrP) [2]. What distinguishes prions is their ability to self-template, converting other copies of the same protein into the prion state. This state is heritable and transmissible, making prions a vector of non-Mendelian inheritance and driving interest in them as epigenetic factors.

Functional prions have since been identified across diverse organisms. In yeast, [ESI+] drives the transgenerational inheritance of an activated chromatin state conferring broad environmental stress resistance [4], and [SMAUG+] allows cells to anticipate nutrient repletion after starvation [5]. The [Het-s] prion of *Podospora anserina* mediates self/non-self recognition [6,7], and the Luminidependens protein of *Arabidopsis thaliana* forms a prion that signals flowering in response to temperature [8]. Although functionally diverse, these proteins share a common feature in the form of prion domains, which confer conformational flexibility, intrinsic disorder, and the capacity to self-template through aggregation. Prion domains share a characteristic composition, combining an enrichment of glutamine (Q) and asparagine (N) residues that supports aggregation through hydrogen bonding [9,10] with a depletion of the charged residues aspartate (D), glutamate (E), lysine (K) and arginine (R) that would otherwise disrupt it [11]. This compositional signature has been exploited by prediction tools such as PLAAC [12], which identify prion-like domains (PrLDs) as candidates for experimental validation.

Combining prediction with experimental confirmation has since identified prions in bacteria [13], archaea [14], and viruses [15], extending them across all three domains of life and beyond cellular life. Computational surveys have identified PrLDs across eukaryotic viruses, including in retroviral Gag [16]. In the nucleocapsid domain of Gag, which packages the viral genome [17], a conserved intrinsically disordered region lies alongside the zinc knuckles and drives zinc-dependent liquid–liquid phase separation that positions and traffics the viral genomic RNA, a feature conserved across retrovirus subfamilies [18]. The breadth of this distribution and the functional diversity of prions have led to the suggestion that they may be an ancient and conserved feature of protein biology.

Investigation of this hypothesis has been difficult. Prion domains are defined by composition rather than by sequence, and the disordered regions that carry them evolve rapidly, which has limited alignment-based evolutionary analysis [19]. Compositional surveys have nonetheless traced prion-like domains across fungal and bacterial lineages [20–22], though how frequently these domains are gained and lost remains unknown. Retrotransposons offer a useful system to answer this. They are mobile genetic elements widespread across eukaryotes, are evolutionary progenitors of retroviruses, undergo similar replication cycles, and share many structural features with retroviral genomic RNA [23]. PrLDs have been described in retroviruses, but no comparable survey has been conducted across retrotransposons, despite functional evidence that these domains matter within them. A PrLD in the N-terminal region of the LTR retrotransposon Ty1, one of the most abundant elements in many laboratory strains of *Saccharomyces cerevisiae* [24,25], is required for retrotransposition and can be replaced by validated yeast prion domains without loss of function [23].

Whether this reflects a general feature of LTR retroelements or a property of Ty1 alone has not been tested. In LTR retroelements, Gag is encoded separately from the conserved Pol proteins, reverse transcriptase (RT), ribonuclease H (RNaseH) and integrase (INT). PrLD presence in Gag can therefore be mapped onto a phylogeny built from sequence that does not contain the domains themselves, avoiding the lack of conservation that has limited evolutionary analysis of disordered regions. We scanned 2,636 proteins from the GyDB for PrLDs, focusing on the six core LTR retroelement proteins Gag, aspartic protease (AP), RT, RNaseH, INT and envelope (ENV) (Fig. 1a), examined their distribution while controlling for disorder, and reconstructed their evolutionary dynamics within Gag. We show that PrLDs are restricted to Gag, that they fall mainly in the C-terminal half of the protein, and that their presence is phylogenetically clustered. They were present in the reconstructed ancestor of these elements and have been lost far more often than gained yet are retained in three of the four superfamilies.

**Figure 1.**
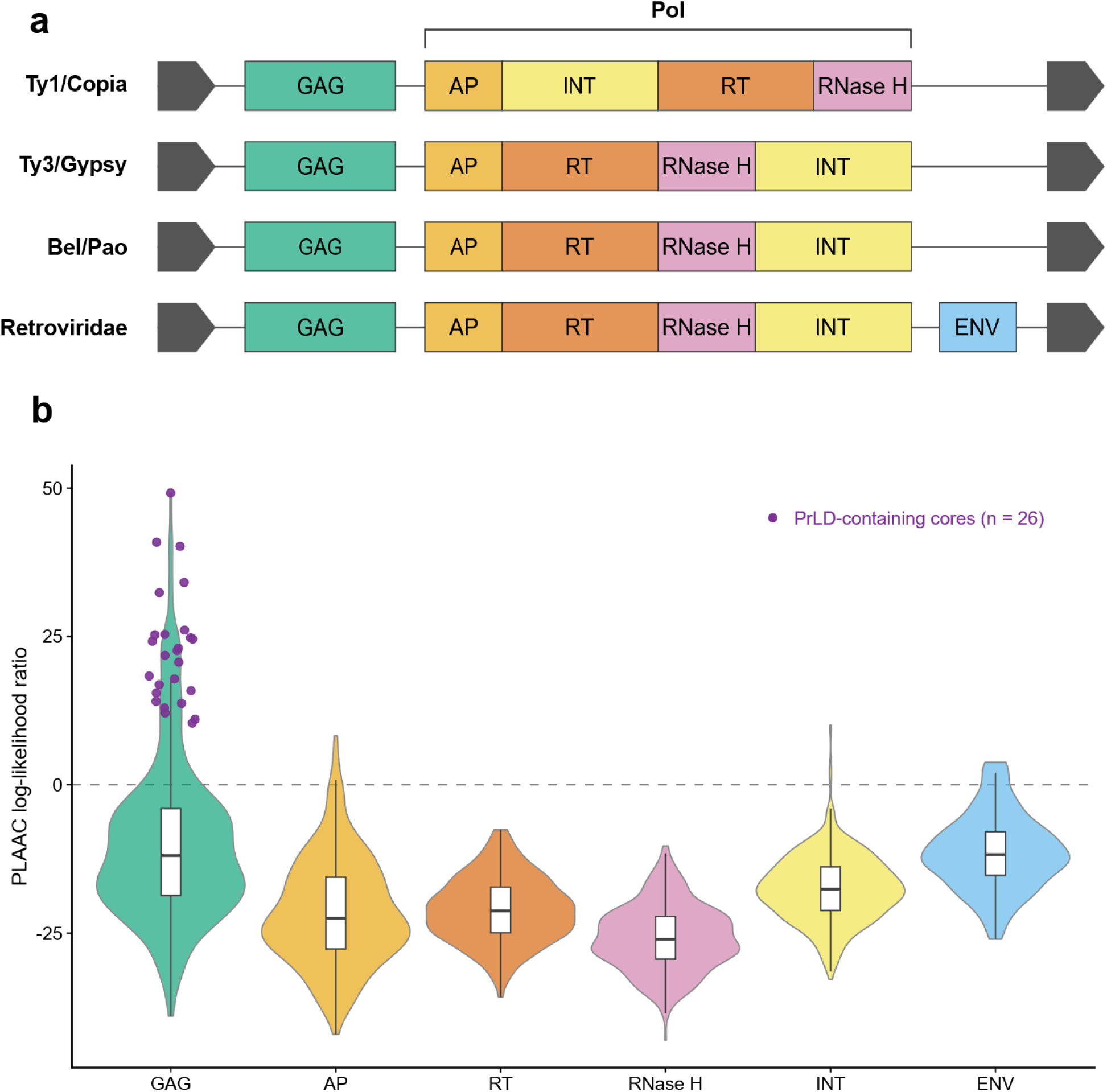
Prion-like domains are restricted to Gag among LTR retroelement proteins. (a) The structure of the four LTR retroelement superfamilies. (b) PLAAC log-likelihood ratio (LLR) for the six core proteins (Gag, n = 331; AP, n = 495; RT, n = 364; RNaseH, n = 368; INT, n = 437; ENV, n = 70). Violins show the full LLR distribution (equal width, not scaled by n); interior boxes show the median and interquartile range. Purple points mark the 26 proteins in which PLAAC called a PrLD, all of which are Gag. The dashed line marks LLR = 0, where positive values indicate more prion-like composition. Thirteen cores with undefined LLR were excluded from this panel.

## Results

### Prion-like domains are restricted to Gag among LTR retroelement proteins

Taking advantage of the strong enrichment of Q/N residues within prion domains, we used PLAAC [12] to scan all 2,636 proteins within the GyDB for PrLDs. PLAAC identified 26 PrLDs, all of which fell in Gag, the only one of 74 protein categories to carry them (26/336 Gag versus 0/2,300 non-Gag; Fisher’s exact test, p < 2.2 × 10⁻¹⁶). Of the 2,300 non-Gag proteins, 1,742 were from the five core LTR retroelement proteins other than Gag (Table S2). Comparing the PLAAC log-likelihood ratios (LLR) of these six core LTR retroelement proteins, Gag and ENV showed comparable median LLR, with Gag distinguished by an extended upper tail driven by the 26 PrLDs rather than an overall shift in distribution (Fig. 1b).

### Gag PrLDs are not explained by disorder alone

Gag contains extensive intrinsically disordered regions (IDRs) [26], so to determine whether PrLD exclusivity within Gag was due to domain-level differences in disorder we identified IDRs in all proteins using metapredict [27] and quantified disorder by measuring the mean disorder of each protein’s highest-scoring IDR of at least 60 aa (Gag 177, non-Gag 253). Proteins containing non-standard amino acids cannot be scored by metapredict and were excluded, as were proteins shorter than 20 residues (n = 146 of the 2,636, including one PrLD-positive protein, GypsySL_monotypic|Chr08_2s54), leaving 2,490 proteins with disorder predictions. Although Gag IDRs overall have higher disorder than non-Gag IDRs (Wilcoxon rank-sum test, p = 2.2 × 10⁻⁴), this difference seems to be largely driven by highly disordered IDRs within PrLD-positive Gag proteins. With these removed, the mean disorder of PrLD-negative Gag and non-Gag IDRs is 0.779 and 0.751 respectively (Wilcoxon rank-sum test, p = 0.042), compared with 0.904 for PrLD-positive Gag, which is significantly higher than PrLD-negative Gag (Wilcoxon rank-sum test, p < 0.001) (Fig. 2a). To control for disorder, proteins were restricted to those whose highest-scoring IDR had a mean disorder of at least the lowest value observed among the PrLD-positive group (0.749) together with the PrLD-positive proteins themselves (n = 257). Within this disorder-matched set, PrLDs remained restricted to Gag (25/126 Gag versus 0/131 non-Gag; Fisher’s exact test, p = 4.7 × 10⁻⁹) (Fig. 2b).

**Figure 2.**
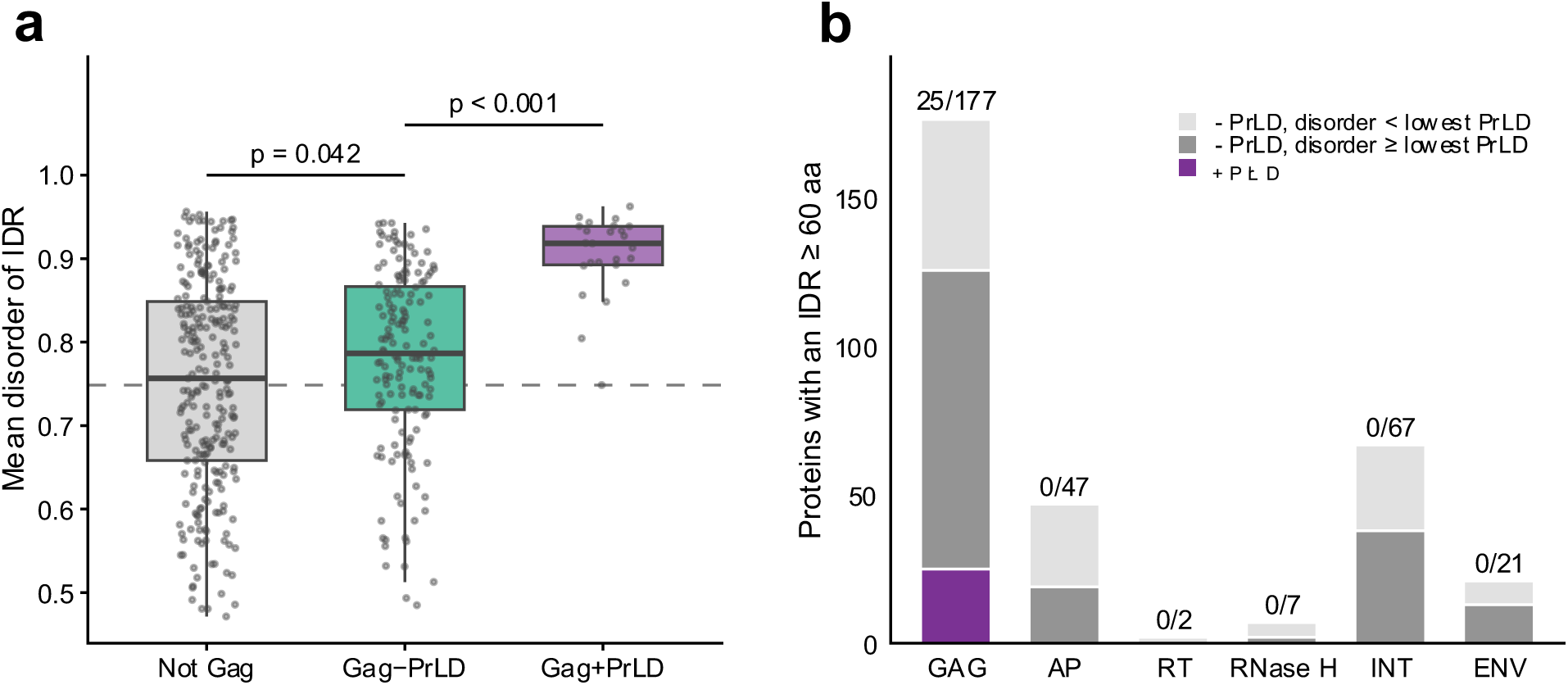
Disorder alone does not explain the restriction of PrLDs to Gag. (a) Mean disorder of each protein’s most disordered IDR of at least 60 residues, for non-Gag (n = 253), PrLD-negative Gag (n = 152) and PrLD-positive Gag (n = 25). Boxes show median and interquartile range; points show individual proteins. The dashed line marks 0.749, the lowest value among PrLD-positive Gag. P values are two-sided Wilcoxon rank-sum tests. (b) Proteins with an IDR of at least 60 residues in each core protein, split by PrLD status and, for PrLD-negative proteins, by whether mean disorder reaches 0.749. Fractions give PrLD-positive out of the total shown. Core proteins only; the disorder-matched test in the text also includes non-core proteins.

### Gag PrLDs share prion-domain composition but are enriched in proline

To determine whether the 26 identified PrLDs were compositionally similar to validated prions but distinct from PrLD-negative Gag IDRs, the fraction of each residue was calculated and compared with that of the yeast prions SUP35, SWI1, CYC8, URE2, MOT3 and RNQ1 [28–30], and with experimentally validated archaeal prion-like domains, in which higher tyrosine (Y) and phenylalanine (F) content distinguished candidates that tested positive in prion assays [14] (Supplementary Fig. S1). As expected, Gag PrLDs, yeast prions and archaeal PrLDs all possessed high Q + N fractions and were depleted in D/E and K/R compared with PrLD-negative IDRs (Fig. 3). Gag PrLDs shared a higher Y + F fraction with archaeal PrLDs than with yeast prions or PrLD-negative Gag IDRs. They were also unusually enriched in proline (P).

**Figure 3.**
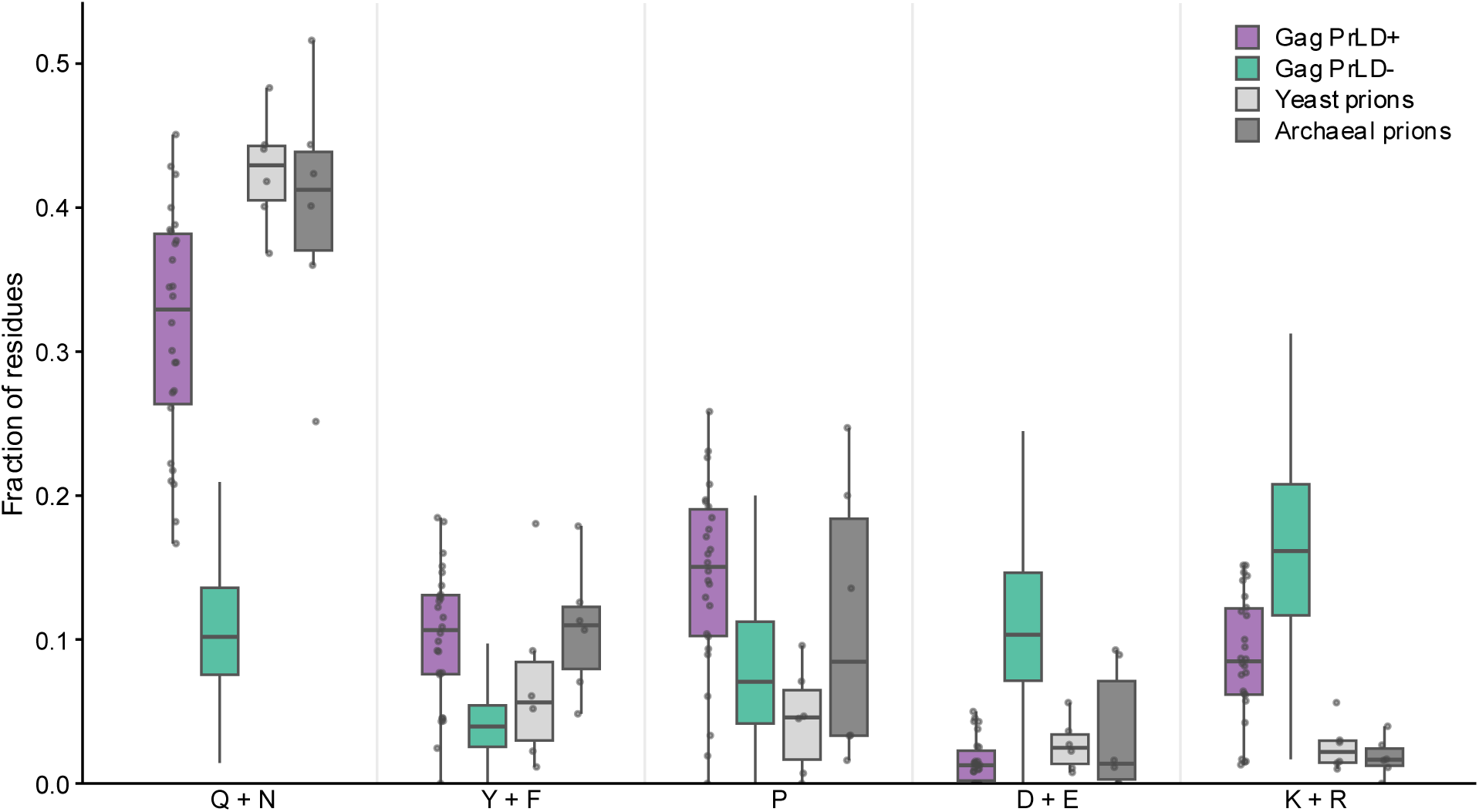
Gag PrLDs share prion-domain composition but are enriched in proline. Fraction of residues in five classes (Q + N, Y + F, P, D + E, K + R) for Gag PrLDs (n = 26), PrLD-negative Gag IDRs (n = 196 IDRs from 152 proteins), yeast prion domains (n = 6) and archaeal prion-like domains (n = 6). Boxes show the median and interquartile range; points show individual domains, omitted for PrLD-negative Gag IDRs for clarity.

### PrLDs are found mainly in the C-terminal half of Gag

Although past work has highlighted the importance of PrLDs and disordered regions in the N-terminus of Gag [23,31], the median relative position of PrLDs was 0.695, with only 3 of 26 falling in the N-terminal half. Two of these were from the Ty1/Copia superfamily (Ty1B, 0.251; Tkm1, 0.323), consistent with findings that an N-terminal PrLD is required for retrotransposition of Ty1 [23]. The third was Cgret (Ty3/Gypsy, 0.389). Within Retroviridae, PrLDs were strongly C-terminal (0.812–0.954). PrLDs had a median length of 79 aa and a range of 60–163 aa (Table S4). To examine the structure of PrLD-positive Gag, Pfam signatures were assigned with InterProScan. Only 14 of 26 PrLD-containing Gag proteins returned Pfam hits. Among these, PrLDs fell downstream of the capsid domain in five of the six proteins with an annotated capsid domain. In all four proteins with an annotated zinc knuckle, the knuckle lay within the PrLD (K-HERV, ASSBSV) or within 11 residues of its end (Cgret, Retrosat-2), placing these PrLDs at the capsid–nucleocapsid junction (Fig. 4).

**Figure 4.**
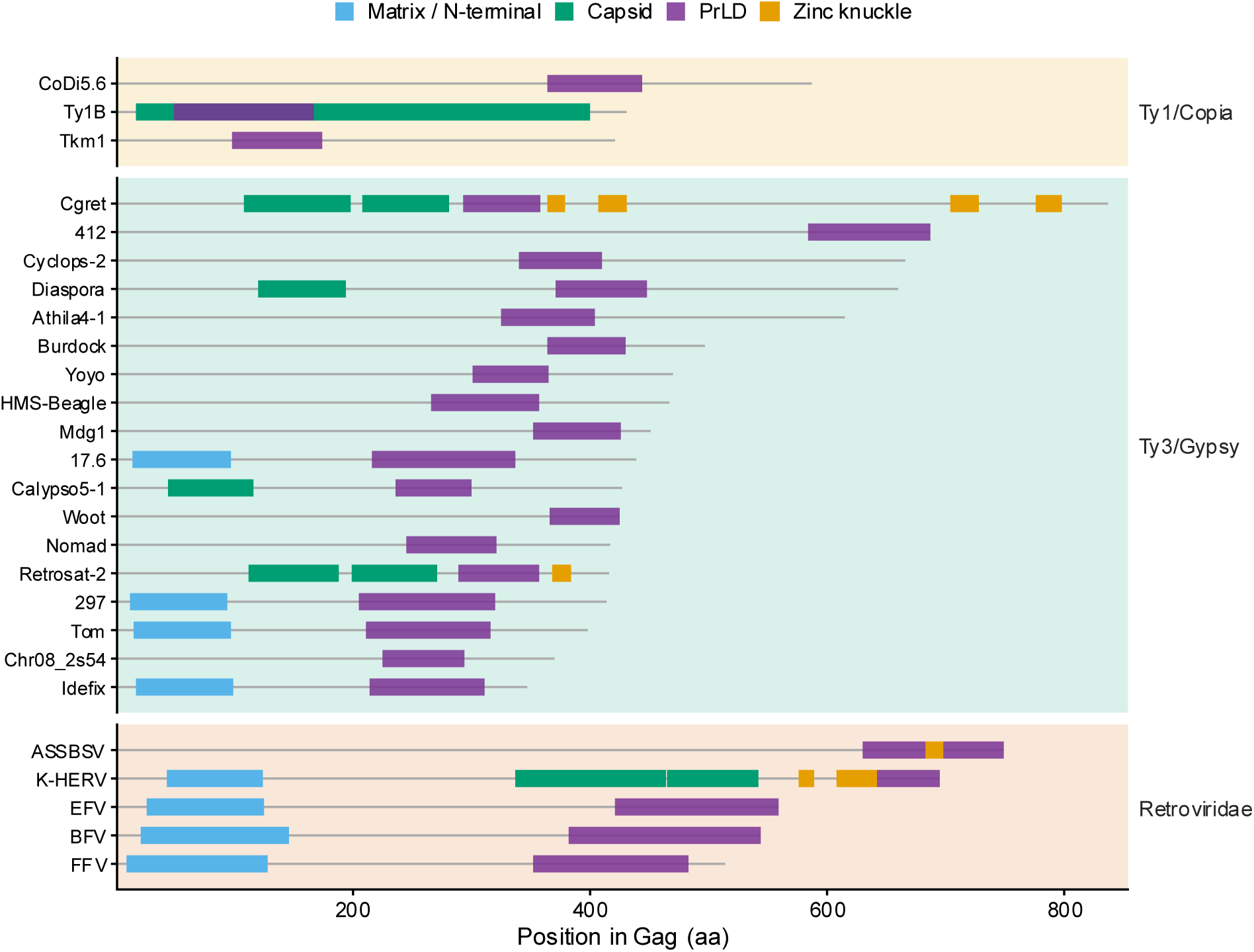
PrLDs occupy the capsid–nucleocapsid region in annotated Gag proteins. Domain organization of the 26 PrLD-containing Gag proteins, grouped by superfamily and ordered by protein length. Grey lines show full protein length; boxes show PrLDs called by PLAAC and Pfam signatures assigned by InterProScan, grouped as matrix/N-terminal, capsid and zinc knuckle (Table S5). Pfam signatures were found in 14 of the 26 proteins.

### PrLD presence is phylogenetically clustered, with recurrent gains and frequent losses

Gag is more variable than the Pol domains and too heterogeneous to align reliably across LTR retroelements [32], so the phylogeny was built from RT, RNaseH and INT and matched to the associated Gag proteins using a shared sequence identifier unique to each element. A maximum-likelihood phylogeny of 258 elements was inferred from 782 amino acid sites, using elements with sequences for all four of Gag, RT, RNaseH and INT (Fig. 5). Elements were excluded if any of the four sequences contained non-standard amino acids. This removed one PrLD-positive element, GypsySL_monotypic|Chr08_2s54, which is also the PrLD-positive protein excluded from the disorder analysis above, so 25 of the 26 PrLDs identified are represented on the tree. The tree was rooted using two methods, with Ty1/Copia as an outgroup and at the midpoint. Under both rootings, all four superfamilies were monophyletic.

**Figure 5.**
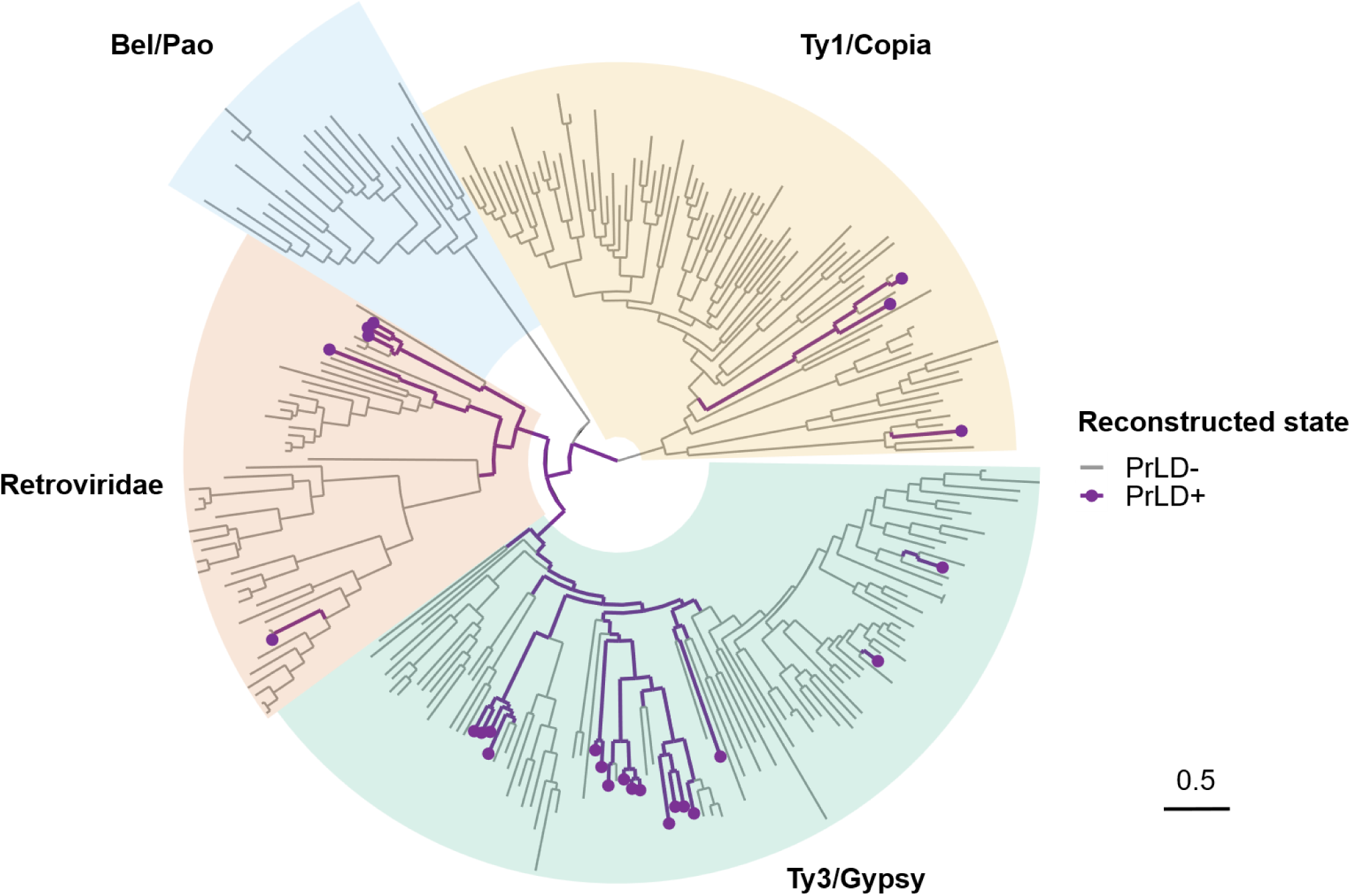
Phylogeny of LTR retroelements with reconstructed PrLD states. Maximum-likelihood tree inferred from concatenated RT, RNaseH and INT sequences (258 elements, 782 sites), rooted on Ty1/Copia. Branches and tips are coloured by PrLD state reconstructed under the ARD model; purple indicates PrLD-positive. Points mark elements in which a PrLD was detected. Superfamily labels give PrLD-positive elements out of the total. Scale bar, 0.5 substitutions per site.

PrLD presence was phylogenetically clustered under both rooting methods (outgroup, D = 0.185; midpoint, D = 0.186; Table 1). When tested against random and Brownian expectations, PrLD patterns differed significantly from random (p < 0.001 under both) but not from Brownian (outgroup, p = 0.288; midpoint, p = 0.298; Supplementary Fig. S2). To reconstruct ancestral PrLD presence within Gag, two models were fitted, an equal-rates (ER) model and an all-rates-different (ARD) model. ER assumes gains and losses occur at equal rates, whereas ARD does not. Under both rootings ARD was the better model, with an AIC 21.6 and 22.2 lower than ER for the outgroup and midpoint rootings respectively (AIC: ER = 155.4 both; ARD = 133.7, outgroup; ARD = 133.2, midpoint; Table 1). This difference was significant (outgroup, p = 1.2 × 10⁻⁶; midpoint, p = 8.7 × 10⁻⁷; Table 1), indicating that gains and losses of PrLDs do not occur at the same rate.

**Table 1.**
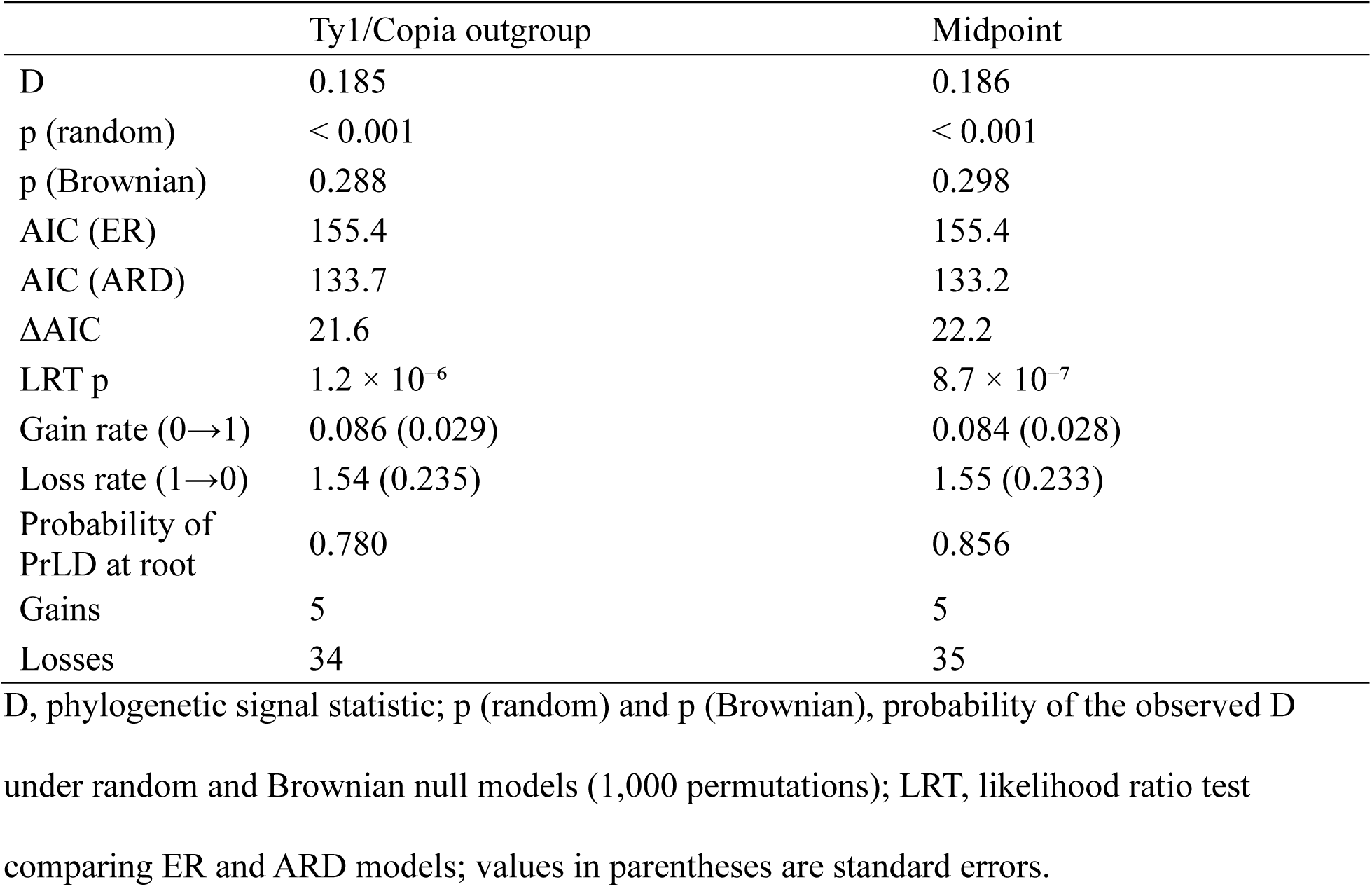
Evolutionary model summary.

The root of the tree was reconstructed as PrLD-positive under both rootings (outgroup, probability = 0.780; midpoint, 0.856). Losses occurred approximately 18 times faster than gains, with gain rates of 0.086 (SE 0.029) under outgroup rooting and 0.084 (SE 0.028) under midpoint rooting, compared with loss rates of 1.54 (SE 0.235) and 1.55 (SE 0.233) respectively. Under this model the absence of PrLDs within Bel/Pao is explained by a single loss at the base of the superfamily (100/100), and in total there were 5 gains and 34 losses under outgroup rooting and 5 gains and 35 losses under midpoint rooting (Supplementary Table S3). Of the predicted gains, three were at internal nodes with high support (99.1/99, 100/100, 100/100) and two were at terminal nodes (Table 2). Of the 34 losses, 12 of the 14 at internal nodes met support thresholds of UFBoot ≥ 95 and SH-aLRT ≥ 80, often affecting whole clades, whereas the losses at node 382 of Retroviridae (88.9/61) and node 508 of Ty3/Gypsy (93/91) did not (Supplementary Table S3). Across the tree, PrLD-positive elements were concentrated in Ty3/Gypsy (17 of 103), with fewer in Retroviridae (5 of 49) and Ty1/Copia (3 of 85), and none in Bel/Pao (0 of 21) (Fig. 5).

**Table 2.** Inferred gains of prion-like domains.

| Gain | Support | Superfamily | Elements | PrLD (+) | Members |
|---|---|---|---|---|---|
| 1 | 99.1/99 | Ty3/Gypsy | 2 | 1 | Cgret, Pyret |
| 2 | Terminal node | Ty3/Gypsy | 1 | 1 | Retrosat-2 |
| 3 | 100/100 | Retroviridae | 2 | 1 | HERV-K10, K-HERV |
| 4 | 100/100 | Ty1/Copia | 5 | 2 | Tkm1, Tse1, Ty1B, Ty2, Ty4 |
| 5 | Terminal node | Ty1/Copia | 1 | 1 | CoDi5.6 |

Support values are SH-aLRT/UFBoot. Terminal node indicates a gain on a terminal branch, for which no support value is defined. PrLD (+) gives the number of PrLD-positive elements among those descending from the gain.

## Discussion

This work indicates that PrLDs within LTR retroelements are a Gag-specific feature, with no PrLDs found among 2,300 non-Gag proteins spanning 73 protein categories. Gag contains extensive disordered regions, but this does not account for the pattern, since the association between PrLDs and Gag remained significant among proteins matched for disorder. Gag is central to multimerisation, RNA binding and virus-like particle assembly in both LTR retrotransposons and retroviruses [23], functions that have also been attributed to PrLDs [33]. A PrLD in the N-terminus of Ty1 Gag is required for retrotransposition, which can be restored by replacing that domain with a validated yeast prion domain [23]. Among the proteins with Pfam annotations, PrLDs lay at the capsid–nucleocapsid junction, adjacent to the nucleic-acid-binding zinc knuckle, placing them in the region of Gag responsible for RNA binding and assembly. The presence and exclusivity of PrLDs in Gag across multiple superfamilies, together with this functional evidence, suggests that they serve a role in Gag rather than being incidental low-complexity sequences.

The 26 Gag PrLDs resemble experimentally validated prion domains but are distinguished from both yeast prion domains and archaeal PrLDs by a high proline fraction. Gag PrLDs showed similar enrichment of Q/N and depletion of D/E and K/R, consistent with prion behaviour, where Q/N drives stable aggregation through hydrogen bonding in cross-β sheets and charged residues destabilise aggregation through charge repulsion [11]. Previous work has suggested a correlation between archaeal PrLDs that tested positive in prion assays and a higher Y/F fraction [14]. Gag PrLDs showed a similar Y/F fraction to these validated archaeal domains. Most notably, Gag PrLDs were unusually enriched in proline, which has previously been considered aggregation-opposing owing to the conformational rigidity imposed by its cyclic side chain [34]. A high proline content has also been reported in the Ty1 Gag PrLD, where it was suggested to prevent secondary structure formation [23]. This enrichment may indicate that Gag PrLDs favour dynamic assembly over stable amyloid formation.

The evidence presented here indicates that PrLDs were present in the Gag of the ancestral LTR retroelement and have since been repeatedly lost and rarely gained. Under an ARD model the root was reconstructed as PrLD-positive under both rooting strategies, and PrLDs were lost approximately 18 times faster than they were gained, with 5 gains and 34 losses. The high loss rate is not surprising, as PLAAC is threshold based and a few substitutions introducing charged residues within a PrLD may bring it below the threshold without necessarily abolishing prion-like behaviour. Consistent with this, the Gag protein of the *Arabidopsis thaliana* element EVD is PLAAC-negative yet forms retrosome-like cytoplasmic granules, and its predicted disordered regions have been proposed to facilitate oligomerization [35]. Despite these limits on detection, the rapid evolution of these domains and the gradual degradation of retroelement sequences, phylogenetic clustering was still observed (D = 0.185), consistent with a conserved role for PrLDs in Gag.

The N-terminal PrLD documented in previous work appears to represent an independent gain within the Ty1/Copia superfamily [23], whereas PrLDs in most other elements fall in the C-terminal half of Gag. This positional difference is consistent with reports that a conserved intrinsically disordered region in the retroviral Gag nucleocapsid regulates zinc-dependent condensate formation and stress granule assembly in HIV-1-expressing cells, repositioning the viral genomic RNA [18]. Both PrLDs identified within Retroviridae that carry annotated zinc fingers showed the same embedded arrangement. The C-terminal bias of PrLDs and their shared ancestry within Ty3/Gypsy are consistent with a common role in these elements, while the PrLD in the Tkm1, Tse1, Ty1B, Ty2, Ty4 clade reflects a single inferred gain retained in two of the five members, with a distinct origin and position. Together, these patterns suggest that prion-like sequence has arisen in Gag more than once and in different regions of the protein.

PLAAC identifies prion-like amino acid composition rather than prion behaviour, so the 26 domains reported here are candidates for experimental testing rather than confirmed prions. The Ty1 Gag PrLD has already been shown to possess prion behaviour and to be interchangeable with validated prion domains [23], so testing these candidates is a reasonable next step. The Gag restriction is defined relative to the protein cores in GyDB, which stores polyproteins as separate domains, so PrLDs located in inter-domain linkers would not have been captured, although the other five core proteins were scanned in full.

Ancestral states were assigned by maximum marginal likelihood without propagating reconstruction uncertainty, so the 5 gains and 34 losses are point estimates. Both rootings recovered nearly identical values, indicating the pattern is not an artefact of root placement. The difference between gain and loss rates may not directly reflect selection. Prion-like composition requires a sustained Q/N-rich, charge-depleted stretch, so a PrLD is far easier to lose than to gain, and an excess of losses is expected even without selection. What stands out is that PrLDs persist at all, retained in elements across three superfamilies despite this bias. Explicit tests of selection on PrLD-containing Gag sequences would determine whether that persistence reflects maintained function.

## Conclusions

This study reveals that prion-like domains are a conserved feature of Gag across LTR retroelements, confined to a single protein among the many these elements encode. All 26 PrLDs identified among 2,636 proteins fell within Gag, and the association held when proteins were matched for intrinsic disorder, indicating that disorder alone does not explain the restriction. These domains share the compositional signature of validated yeast and archaeal prion domains but carry an unusual proline enrichment, suggesting they may favour dynamic assembly over stable amyloid. Mapping PrLD presence onto a phylogeny built from the conserved Pol proteins shows that the domains were present in the ancestor of these elements and have been lost far more often than gained, yet retained across three superfamilies, indicating maintenance rather than repeated acquisition. Collectively, these findings establish PrLDs as a general feature of Gag rather than incidental low-complexity sequence, provide a route to reconstructing the history of compositionally defined domains in lineages where they cannot be aligned, and identify 26 candidates for experimental testing.

## Methods

### Dataset and domain annotation

Protein sequences were obtained from the CORES database of GyDB 2.0 [36] (accessed 2026-08-08). The database contains 2,636 protein cores from mobile genetic elements, referred to throughout as proteins. Each sequence label begins with the protein’s category (e.g., GAG, RT), which was used to assign proteins to groups. These include the six core proteins of LTR retroelements (LTR retrotransposons and retroviruses): Gag, AP, RT, RNaseH, INT and ENV. The remainder are accessory ORFs and domains from other mobile genetic elements. The mean lengths of the main LTR retroelement proteins were 360 aa (Gag), 472 aa (ENV), 327 aa (INT), 246 aa (RT), 154 aa (AP) and 142 aa (RNaseH) (Table S1).

### Identification of Prion-like domains

Prion-like domains were predicted with PLAAC using the command-line version (v1.1.0b7; https://github.com/whitehead/plaac) [12]. PLAAC was run on all 2,636 GyDB proteins with default parameters (minimum PrLD length of 60 amino acids), except for α = 0.5, following the developers’ approach for non-yeast proteins [12]. This weights the background amino-acid frequencies equally between *Saccharomyces cerevisiae* and the GyDB core set, balancing yeast-specific and input-specific compositional bias. Proteins were classified as PrLD-positive when PLAAC reported a PrLD length greater than zero. To test whether PrLDs were enriched in Gag, the proportion of Gag proteins containing a PrLD was compared with that of non-Gag proteins using Fisher’s exact test.

### Prediction of intrinsically disordered regions

Metapredict (v2.65.1) [27] was used to identify IDRs within GyDB proteins. Proteins containing non-standard amino acids cannot be scored by metapredict and were excluded, as were proteins shorter than 20 residues (n = 146 of the 2,636, including one PrLD-positive protein). For each remaining protein, the mean disorder of every IDR of at least 60 residues was calculated using the boundaries and per-residue scores reported by metapredict, and the highest of these values was retained. The 60-residue threshold matches PLAAC’s minimum PrLD length and restricts the analysis to proteins with IDRs long enough for PLAAC to identify a PrLD. Proteins lacking PrLDs were classified as similarly disordered if their mean disorder was higher than or equal to the lowest value observed among PrLD-positive proteins.

To assess the impact of disorder on PrLD detection, mean disorder scores were compared using two-sided Wilcoxon rank-sum tests for Gag versus non-Gag, PrLD-positive Gag versus PrLD-negative Gag, and PrLD-negative Gag versus non-Gag. To determine whether the enrichment of PrLDs in Gag proteins could be explained by differences in disorder, the proportion of proteins containing a PrLD was compared between Gag and non-Gag proteins within the disorder-matched set, comprising PrLD-positive proteins together with those whose highest-scoring IDR reached the lowest value observed among PrLD-positive proteins, using Fisher’s exact test. Comparisons by protein category were restricted to the six core LTR retroelement proteins.

### Analysis of Amino acid composition of the PrLDs

The composition of Gag PrLDs was compared with validated yeast prion domains, experimentally validated archaeal prion-like domains, and PrLD-negative Gag IDRs. Gag PrLD sequences were taken from the prion domain sequence (PRDaa) reported in the PLAAC output. PrLD-negative Gag IDR sequences were extracted from the GyDB proteins using the metapredict boundaries described above, retaining IDRs of at least 60 residues. The experimentally validated yeast prions SUP35, SWI1, CYC8, URE2, MOT3, and RNQ1 were selected from the literature for being Q/N-rich [28–30] to compare with the Q/N-rich PrLDs identified with PLAAC. Yeast prion amino acid sequences were obtained from UniProt, namely P05453 (SUP35), P09547 (SWI1), P14922 (CYC8), P23202 (URE2), P54785 (MOT3) and P25367 (RNQ1) [37]. PLAAC was run on all six proteins with default parameters, which weight the background toward *Saccharomyces cerevisiae* and are appropriate for yeast sequences, and PrLDs were taken from the PRDaa field of the PLAAC output. Archaeal PrLDs were obtained from Dataset S3 of Zajkowski et al. [14], restricted to candidates scoring positive in both the Congo red binding and yeast prion reporter assays (n = 6). For each domain, the fraction of residues in five classes (Q + N, Y + F, P, D + E, K + R) was calculated and compared between groups (Fig. 3).

### Determining where PrLDs are located within Gag

Gag proteins were annotated with Pfam signatures from InterProScan [38] via the EMBL-EBI web service (run 13 August 2026). Pfam accessions were grouped into three classes, matrix/N-terminal, capsid and zinc knuckle (Table S5). Annotated domains were recovered for 14 of 26 PrLD-containing Gag proteins. PrLD position was expressed as the midpoint of the PrLD divided by protein length, with values below 0.5 indicating the N-terminal half and values higher than 0.5 indicating the C-terminal half.

### Phylogenetic signal and ancestral reconstruction

The phylogenetic tree was built from RT, RNaseH, and INT sequences. Only elements that possessed sequences for all 3 proteins and a Gag protein were included and were separated into protein domain specific fasta files. Sequences with non-standard amino acids were excluded. Fasta files were aligned individually with MAFFT (v7.525) [39] using FFT-NS-i selected from the automatic model selection parameter. Columns with gaps in more than 70% of sequences were removed with trimAl (v1.5.1) [40], and the three trimmed alignments were concatenated by element (258 sequences and 782 amino acid sites). A maximum-likelihood tree was inferred with IQ-TREE v3.1.3 (random seed 12345) [41] selecting the best-fitting substitution model, Q.PFAM+F+R8, by Bayesian information criterion (BIC) with ModelFinder [42] from LG, WAG, JTT, VT, rtREV, HIVb, HIVw, Q.pfam, Dayhoff, Blosum62. Branch support was assessed with 1,000 ultrafast bootstrap replicates, optimized by NNI [43], and 1,000 SH-aLRT replicates [44]. Branches were considered well supported at UFBoot ≥ 95 and SH-aLRT ≥ 80.

The tree was rooted and compared using two approaches. Ty1/Copia is conventionally used as the outgroup for LTR retroelement phylogenies [32,36]. To test whether the results depended on root placement, the tree was rooted both with Ty1/Copia as the outgroup, using root in ape (v5.8.1) [45], and at the midpoint of the longest tip-to-tip path, using midpoint in phangorn (v2.12.1) [46]. To determine the phylogenetic signal, PrLD presence was coded as a binary trait across the 258 tips and quantified with the D statistic for both trees [47] using phylo.d in caper (v1.0.4) [48], with 1,000 permutations and random seed 1234. phylo.d compares the observed D against two models, random association of the trait across the tree, and the clumping expected under a Brownian threshold model.

Ancestral states were reconstructed by maximum likelihood with ace in ape [45], treating PrLD presence as a binary character. Two models were fitted, an ER model and an ARD model. ER assumes gains and losses occur at the same rate whereas ARD allows them to differ. The two models were then compared by Akaike information criterion (AIC) and by a likelihood ratio test. Under the better-fitting model, each internal node was assigned PrLD-positive when the marginal likelihood was greater than 0.5, and gains and losses were counted as transitions between parent and child states along each branch. Branch support for each inferred transition on the outgroup-rooted tree was taken from the SH-aLRT and UFBoot values of the corresponding node, applying the same thresholds as above, and each transition was assigned to a superfamily from the elements descending from it. Rooting splits a single branch into two, leaving one node adjacent to the root with no support label. Support for this node was therefore taken from the sibling node (100/100). All analyses were repeated on both the outgroup-rooted and midpoint-rooted trees for comparison.

## Declarations

### Ethics approval and consent to participate

Not applicable

### Consent for publication

Not applicable

### Availability of data and materials

All data and analysis scripts are available at: https://github.com/CameronDebellefeuille/Retroelement-Prion-Phylogeny

### Authors’ contributions

T.Z. conceived the study, supervised the project, and provided feedback on the manuscript. C.D. designed the study, performed the data analysis, and wrote the manuscript. Both authors read and approved the final version of the manuscript.

## Acknowledgements

We thank the Prioneers research group of the Blue Marble Space Young Scientist Program for discussion and feedback throughout this work, and Blue Marble Space for organizational support.

## Additional files

**Additional file 1:** Supplementary Material. Figures S1–S2 and Tables S1–S5.

