## Supplementary Material (Figures S1-S2, Tables S1-S5) for "Evolutionary dynamics of Gag-restricted prion-like domains in LTR retroelements"

**Contents list:**

Figure S1. Amino acid composition of prion-like domains and reference sets

Figure S2. D statistic null distributions under both rootings

Table S1. Length of GyDB 2.0 LTR protein cores

Table S2. PrLD presence among all GyDB cores

Table S3. All inferred transitions

Table S4. Prion-like domains identified in Gag proteins

Table S5. Pfam signatures used to annotate Gag domain architecture

**Figure S1. Amino acid composition of prion-like domains and reference sets**

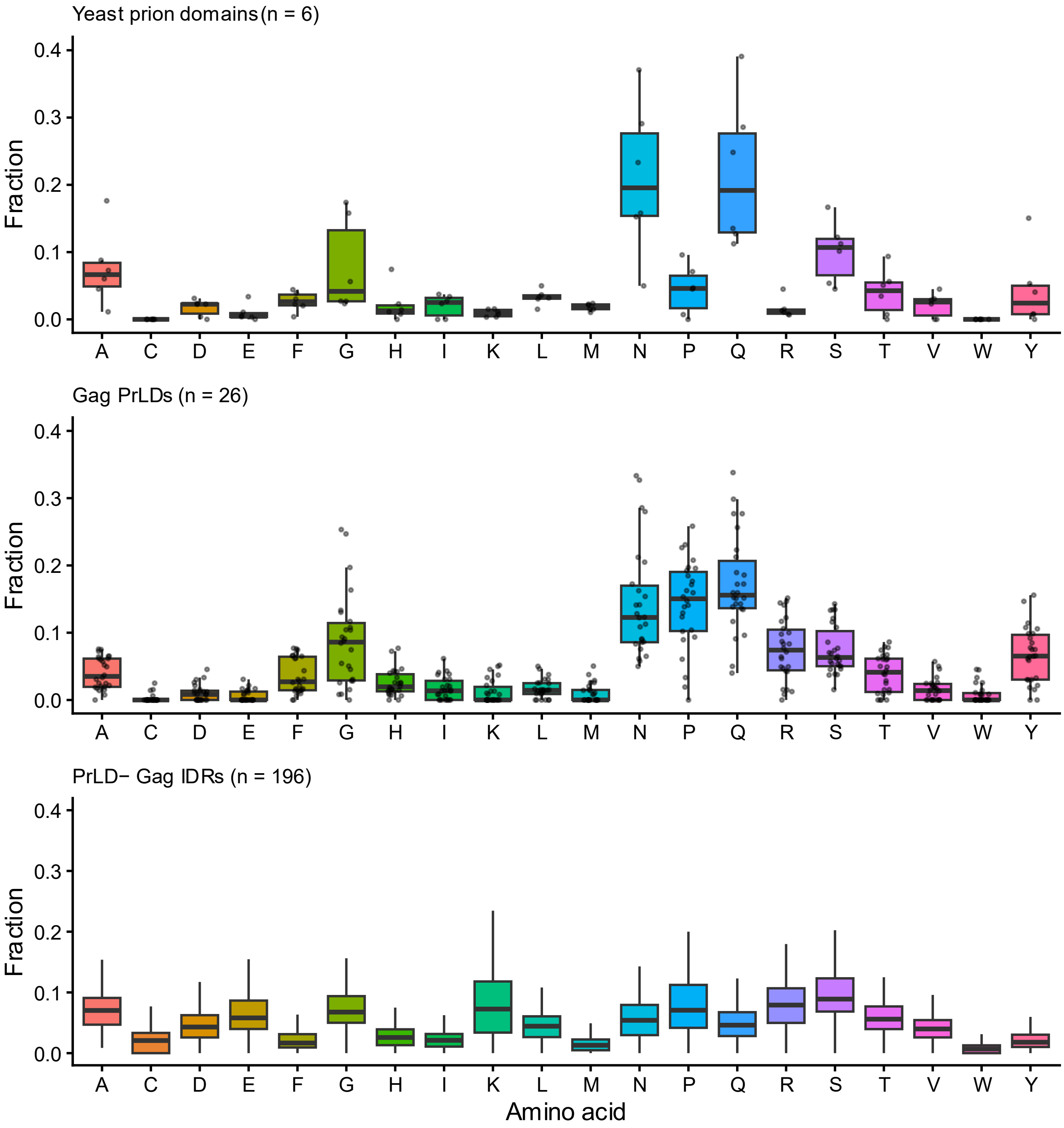

**Figure S2. D statistic null distributions under both rootings**

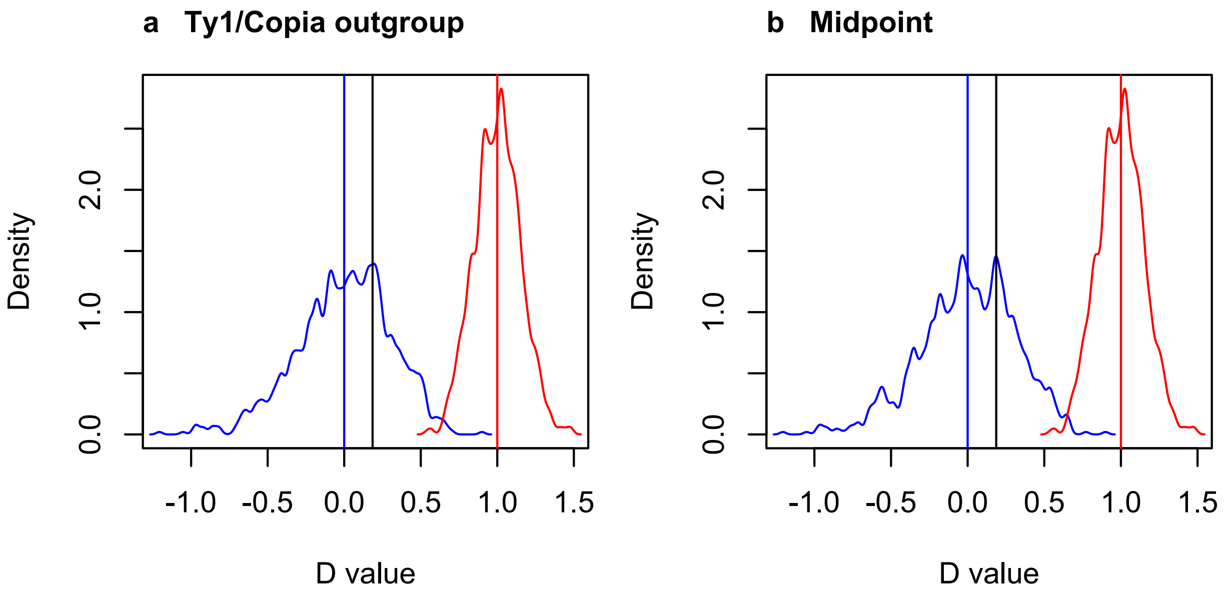

**Table S1. Length of GyDB 2.0 LTR protein cores**

| Protein | Number of cores | Mean length (aa) | Median length (aa) | Range (aa) |
|---|---|---|---|---|
| Gag | 336 | 360.0 | 323.0 | 25–853 |
| AP | 500 | 153.6 | 116.0 | 44–986 |
| RT | 365 | 246.0 | 237.0 | 54–438 |
| RNaseH | 370 | 141.7 | 129.5 | 22–508 |
| INT | 437 | 327.4 | 302.0 | 112–1,122 |
| ENV | 70 | 472.1 | 405.5 | 111–1,225 |

AP, aspartic protease; RT, reverse transcriptase; RNaseH, ribonuclease H; INT, integrase; ENV, envelope. Cores as defined by GyDB 2.0 (Llorens et al., 2011).

**Table S2. PrLD presence among all GyDB cores**

| Protein | PrLD+ | PrLD− | Total |
|---|---|---|---|
| **Gag** | **26** | **310** | **336** |
| AP | 0 | 500 | 500 |
| RT | 0 | 365 | 365 |
| RNaseH | 0 | 370 | 370 |
| INT | 0 | 437 | 437 |
| ENV | 0 | 70 | 70 |
| Other GyDB | 0 | 558 | 558 |
| **Total non-Gag** | **0** | **2,300** | **2,300** |

AP, aspartic protease; RT, reverse transcriptase; RNaseH, ribonuclease H; INT, integrase; ENV, envelope. Cores as defined by GyDB 2.0 (Llorens et al., 2011).

**Table S3. All inferred transitions**

| Change | Label | Type | Support | Superfamily | Elements |
|---|---|---|---|---|---|
| gain | node_413 | internal | 100/100 | Retroviridae | 2 |
| gain | node_337 | internal | 100/100 | Ty1/Copia | 5 |
| gain | CoDi5.6 | terminal | NA | Ty1/Copia | 1 |
| gain | node_473 | internal | 99.1/99 | Ty3/Gypsy | 2 |
| gain | Retrosat-2 | terminal | NA | Ty3/Gypsy | 1 |
| loss | node_357 | internal | 100/100 | Bel/Pao | 21 |
| loss | ZFERV-2 | terminal | NA | Retroviridae | 1 |
| loss | node_382 | internal | 88.9/61 | Retroviridae | 13 |
| loss | SnRV | terminal | NA | Retroviridae | 1 |
| loss | node_395 | internal | 100/100 | Retroviridae | 26 |
| loss | HERV-K10 | terminal | NA | Retroviridae | 1 |
| loss | MuERV-L | terminal | NA | Retroviridae | 1 |
| loss | node_424 | internal | 100/100 | Retroviridae | 2 |
| loss | DrFV-1 | terminal | NA | Retroviridae | 1 |
| loss | node_515 | internal | 100/100 | Ty1/Copia | 85 |
| loss | Tse1 | terminal | NA | Ty1/Copia | 1 |
| loss | Ty2 | terminal | NA | Ty1/Copia | 1 |
| loss | Ty4 | terminal | NA | Ty1/Copia | 1 |
| loss | Tv1 | terminal | NA | Ty3/Gypsy | 1 |
| loss | node_508 | internal | 93/91 | Ty3/Gypsy | 3 |
| loss | Ted | terminal | NA | Ty3/Gypsy | 1 |
| loss | Zam | terminal | NA | Ty3/Gypsy | 1 |
| loss | Tor4a | terminal | NA | Ty3/Gypsy | 1 |
| loss | node_503 | internal | 100/100 | Ty3/Gypsy | 2 |
| loss | node_454 | internal | 99.6/98 | Ty3/Gypsy | 45 |
| loss | Pyret | terminal | NA | Ty3/Gypsy | 1 |
| loss | node_498 | internal | 100/100 | Ty3/Gypsy | 2 |
| loss | Cigr-1 | terminal | NA | Ty3/Gypsy | 1 |
| loss | node_501 | internal | 99.6/100 | Ty3/Gypsy | 2 |
| loss | Osvaldo | terminal | NA | Ty3/Gypsy | 1 |
| loss | Bagy-2 | terminal | NA | Ty3/Gypsy | 1 |
| loss | GypsySL_04 | terminal | NA | Ty3/Gypsy | 1 |
| loss | node_445 | internal | 100/100 | Ty3/Gypsy | 8 |
| loss | node_436 | internal | 100/100 | Ty3/Gypsy | 3 |
| loss | node_438 | internal | 100/100 | Ty3/Gypsy | 2 |

| | | | | | |
|---|---|---|---|---|---|
| loss | node_426 | internal | 100/100 | Ty3/Gypsy | 10 |
| loss | Cer1 | terminal | NA | Ty3/Gypsy | 1 |
| loss | Tor1 | terminal | NA | Ty3/Gypsy | 1 |
| loss | Tor2 | terminal | NA | Ty3/Gypsy | 1 |

**Table S4. Prion-like domains identified in Gag proteins**

| Element | Superfamily | Length (aa) | PrLD start | PrLD end | PrLD length | Relative |
|---|---|---|---|---|---|---|
| Ty1B | Ty1/Copia | 431 | 49 | 167 | 119 | 0.251 |
| Tkm1 | Ty1/Copia | 421 | 98 | 174 | 77 | 0.323 |
| CoDi5.6 | Ty1/Copia | 587 | 364 | 444 | 81 | 0.688 |
| Cgret | Ty3/Gypsy | 837 | 293 | 358 | 66 | 0.389 |
| Cyclops-2 | Ty3/Gypsy | 666 | 340 | 410 | 71 | 0.563 |
| Athila4-1 | Ty3/Gypsy | 615 | 325 | 404 | 80 | 0.593 |
| Diaspora | Ty3/Gypsy | 660 | 371 | 448 | 78 | 0.62 |
| Calypso5-1 | Ty3/Gypsy | 427 | 236 | 300 | 65 | 0.628 |
| 17.6 | Ty3/Gypsy | 439 | 216 | 337 | 122 | 0.63 |
| 297 | Ty3/Gypsy | 414 | 205 | 320 | 116 | 0.634 |
| Tom | Ty3/Gypsy | 398 | 211 | 316 | 106 | 0.662 |
| HMS-Beagle | Ty3/Gypsy | 467 | 266 | 357 | 92 | 0.667 |
| Nomad | Ty3/Gypsy | 417 | 245 | 321 | 77 | 0.679 |
| Chr08_2s54 | Ty3/Gypsy | 370 | 225 | 294 | 70 | 0.701 |
| Yoyo | Ty3/Gypsy | 470 | 301 | 365 | 65 | 0.709 |
| Idefix | Ty3/Gypsy | 347 | 214 | 311 | 98 | 0.756 |
| Retrosat-2 | Ty3/Gypsy | 416 | 289 | 357 | 69 | 0.776 |
| Burdock | Ty3/Gypsy | 497 | 364 | 430 | 67 | 0.799 |
| Mdg1 | Ty3/Gypsy | 451 | 352 | 426 | 75 | 0.863 |
| 412 | Ty3/Gypsy | 687 | 584 | 687 | 104 | 0.925 |
| Woot | Ty3/Gypsy | 425 | 366 | 425 | 60 | 0.931 |
| FFV | Retroviridae | 514 | 352 | 483 | 132 | 0.812 |
| BFV | Retroviridae | 544 | 382 | 544 | 163 | 0.851 |
| EFV | Retroviridae | 559 | 421 | 559 | 139 | 0.877 |
| ASSBSV | Retroviridae | 749 | 630 | 749 | 120 | 0.921 |
| K-HERV | Retroviridae | 695 | 631 | 695 | 65 | 0.954 |

Chr08_2s54 (full label GypsySL_monotypic|Chr08_2s54)

**Table S5. Pfam signatures used to annotate Gag domain**

| Accession | Description | Class |
|---|---|---|
| PF02337 | Retroviral GAG p10 protein | Matrix / N-terminal |
| PF03276 | Spumavirus gag protein, N-terminal | Matrix / N-terminal |
| PF30021 | Retroviral Gag protein N-terminal domain | Matrix / N-terminal |
| PF00607 | gag protein p24 N-terminal domain | Capsid |
| PF01021 | Ty transposon capsid protein | Capsid |
| PF03732 | Retrotransposon-derived protein PEG10, N-terminal capsid-like domain | Capsid |
| PF19317 | Gag protein p24 C-terminal domain | Capsid |
| PF30901 | PEG10/RTL1, C-terminal capsid-like domain | Capsid |
| PF00098 | Zinc knuckle | Zinc knuckle |
| PF03716 | WCCH motif | Zinc knuckle |
| PF14787 | GAG-polyprotein viral zinc-finger | Zinc knuckle |
